# External introductions helped drive and sustain the high incidence of HIV-1 in rural KwaZulu-Natal, South Africa

**DOI:** 10.1101/119826

**Authors:** David A. Rasmussen, Eduan Wilkinson, Alain Vandormael, Frank Tanser, Deenan Pillay, Tanja Stadler, Tulio de Oliveira

## Abstract

Despite increasing access to antiretroviral therapy, HIV incidence in rural KwaZulu-Natal communities remains among the highest ever reported in Africa. While many epidemiological factors have been invoked to explain this high incidence, widespread human mobility and movement of viral lineages between geographic locations have implicated high rates of transmission across communities. High rates of cross-community transmission call into question how effective increasing local coverage of antiretroviral therapy will be at preventing new infections, especially if many new cases arise from external introductions. To help address this question, we use a new phylodynamic modeling approach to estimate both changes in epidemic dynamics through time and the relative contribution of local transmission versus external introductions to overall incidence from HIV-1 subtype C phylogenies. Our phylodynamic estimates of HIV prevalence and incidence are remarkably consistent with population-based surveillance data. Our analysis also reveals that early epidemic dynamics in this population were largely driven by a wave of external introductions. More recently, we estimate that anywhere between 20-60% of all new infections arise from external introductions from outside the local community. These results highlight the power of using phylodynamic methods to study generalized HIV epidemics and the growing need to consider larger-scale regional transmission dynamics above the level of local communities when designing and testing prevention strategies.

## Introduction

While the HIV epidemic hit South Africa relatively late compared with other southern African nations, the epidemic grew explosively in the 1990’s from an estimated 0.8% prevalence in 1990 to over 20% in 2000 [UNAIDS, 2014]. Prevalence nationwide stabilized at around 20% in 2000, but an estimated 6.19 million people are still currently living with HIV in South Africa, more than any other country in the world. Prevalence is highest in the province of KwaZulu-Natal (KZN) where it is as high as 25-40% in some communities [Kharsany et al., 2015, Shisana et al., 2015]. While the increasing lifespan of infected individuals on antiretroviral treatment (ART) can partly explain the continued high prevalence, incidence also remains alarmingly high at between 3 and 6% per year in KZN [Karim et al., 2011, Nel et al., 2012], and may be even higher in certain high risk groups such as young women [de Oliveira et al., 2017].

Many factors have been implicated in the explosive growth of the HIV epidemic in southern Africa and KZN in particular, but patterns of human movement in the region have long received special attention [Jochelson et al., 1991, Quinn, 1994, Lurie et al., 1997]. Migration rates have historically been high in the region, fueled by labor-intensive industries such as mining [Hargrove, 2008, Corno and De Walque, 2012]. Mobility has also increased significantly since the end of Apartheid, and sexual networks are known to be geographically well-connected across long distances [Harrison et al., 2008]. However, the exact role human movement has played in the dynamics of the epidemic has been debated [Coffee et al., 2007]. Given the rural and rather isolated nature of many KZN communities, it is apparent that some spatial mixing would have been necessary to seed the epidemic in different geographic locations. Beyond seeding local epidemics, increasing mobility and migrant labor may have also played a larger role by placing individuals at an increased risk of infection [Lurie et al., 2003, Welz et al., 2007, Camlin et al., 2010], possibly due to migrants engaging in riskier sexual behavior outside of their home communities [Coffee et al., 2007].

Resolving the contribution of human movement to the HIV epidemic has been challenging due to the difficulty of quantifying the extent to which transmission occurs locally within communities versus new cases being imported through external introductions. While the geographic source of new infections cannot typically be resolved using traditional surveillance data, viral phylogenies can help reveal the source of new infections [Holmes, 2004, Pybus and Rambaut, 2009, Faria et al., 2011]. Broadly speaking, if new infections primarily arise from transmission within local communities, viral samples collected within the community should be more closely phylogenetically related to one another than to samples taken from outside the community; whereas if many new infections are being externally introduced then local samples will tend to cluster with external samples throughout the tree. Phylogenetics can therefore help reveal the movement of viral lineages within and between different communities. For example, one recent study revealed that a high proportion of new infections in a rural Ugandan community arose from external introductions [Grabowski et al., 2014]. Recent phylogenetic studies have also revealed widespread viral movement across larger spatial scales and even across national borders [Gray et al., 2009, Wilkinson et al., 2016].

Here we explore the role of external introductions in the Africa Health Research Institute’s study population in rural KwaZulu-Natal. Using a phylodynamic approach that couples phylogenetic methods together with epidemiological modeling, we reconstructed epidemic dynamics that are remarkably consistent with population-based surveillance data. By tracking the movement of viral lineages between populations, we also directly quantified the contribution of local transmission versus external introductions to overall HIV incidence. We found that far from just seeding the local epidemic, external introductions played a large role in sustaining high HIV incidence, thus confirming the important role human mobility and migration have played in this hyper-epidemic setting in KZN.

## Results

To help situate the local epidemic in the Africa Health Research Institute (AHRI) study population within the larger context of the southern African HIV epidemic, a maximum likelihood (ML) phylogeny was reconstructed from HIV-1 subtype C sequences sampled from 1,068 infected individuals in the AHRI population along with 11,289 sequences from a larger regional background dataset [Wilkinson et al., 2016] sampled throughout southern Africa (Fig 1 A). Branch lengths in the ML phylogeny were then rescaled into units of calendar time using least squares dating. Although there are some larger clades composed predominantly of local viral samples which likely represent locally evolving sub-epidemics, the majority of samples from the AHRI are interspersed throughout clades composed predominantly of external samples (i.e. from the regional background dataset), suggesting that many independent introduction events have occurred into the local population.

**Figure 1.**
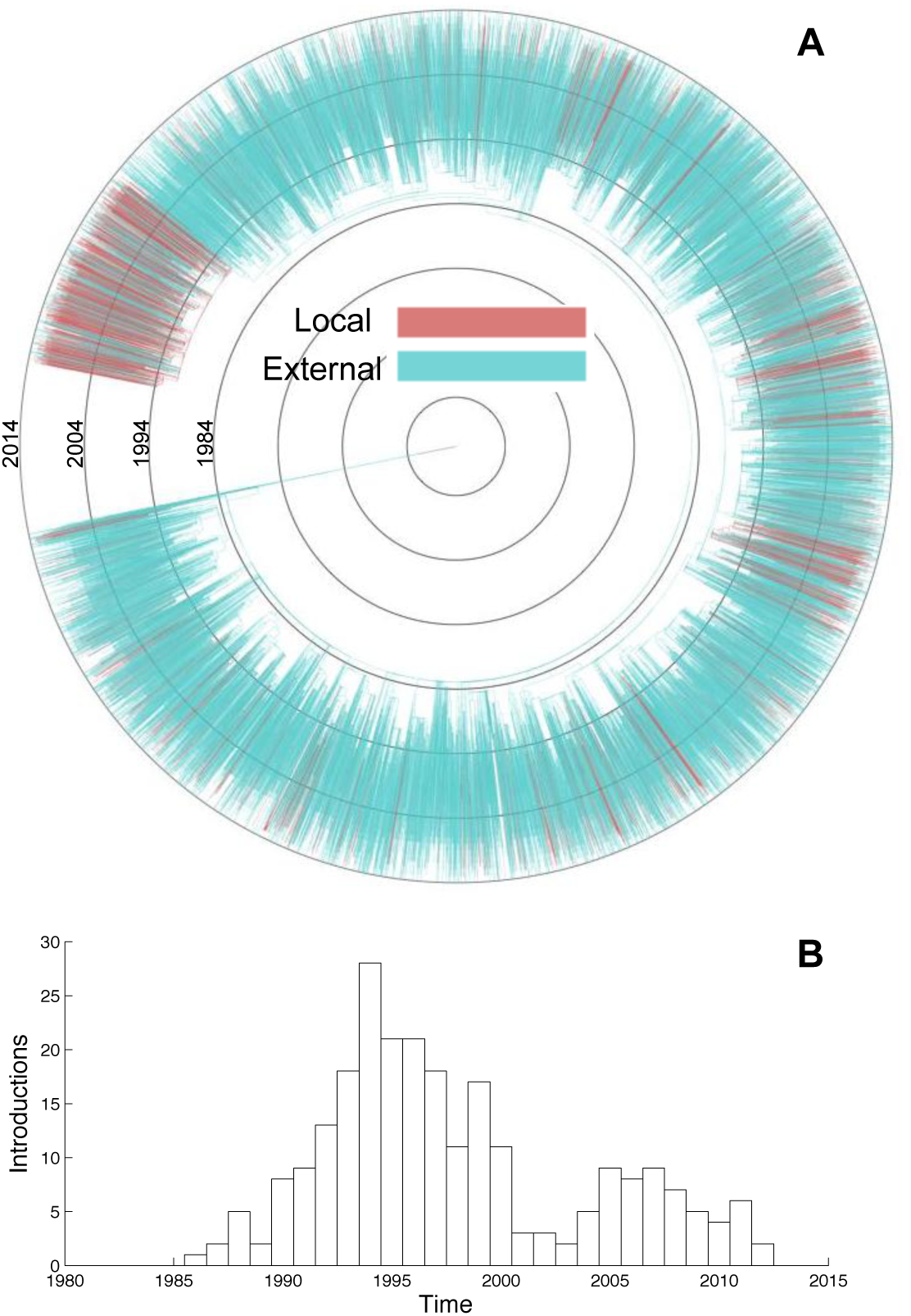
The local HIV epidemic in the AHRI study population within the larger phylogenetic context of the southern African subtype C epidemic. **(A)** Maximum likelihood phylogenetic tree reconstructed from HIV *pol* sequences from the AHRI (local) along with the regional background dataset (external). Tips are colored by sampling location, internal branches are colored according to their ancestral location reconstructed via maximum parsimony. **(B)** Time series showing the temporal distribution of external introductions into the local population identified by maximum parsimony.

The ancestral location of each lineage in the ML tree was then reconstructed using maximum parsimony to reveal how viral lineages have moved over time (Fig 1A). This allowed us to identify potential external introductions at branches in the phylogeny where the most parsimonious ancestral location transitioned from the external to the local population. In total, 248 external introductions were identified into the local population. Using the node heights at which transitions in ancestral locations occurred as a proxy for the timing of external introductions suggested that most of these events occurred between 1990 and 2000 during the period of rapid epidemic growth in South Africa, with relatively fewer introductions after 2000 once the epidemic stabilized (Fig 1B).

Because phylogenies only contain lineages ancestral to sampled viruses, the number of external introductions identified in the foregoing analysis likely represents only a small fraction of the total number. Moreover, due to the large size of the epidemic relative to the number of infected individuals sampled (∼8%), it is extremely unlikely that we would have sampled both descendants of any recent transmission event between an external donor and a local recipient. In fact, the most recent common ancestor of most pairs of sampled viruses predates the early stages of the South African epidemic (Fig 1A). The heights of these nodes may therefore not be a reliable proxy for the timing of introduction events since introductions may have occurred more recently (see Fig 2C). It is highly likely then that our preliminary analysis based on parsimony both underestimates the true number of external introductions and skews their temporal distribution towards the more distance past.

**Figure 2.**
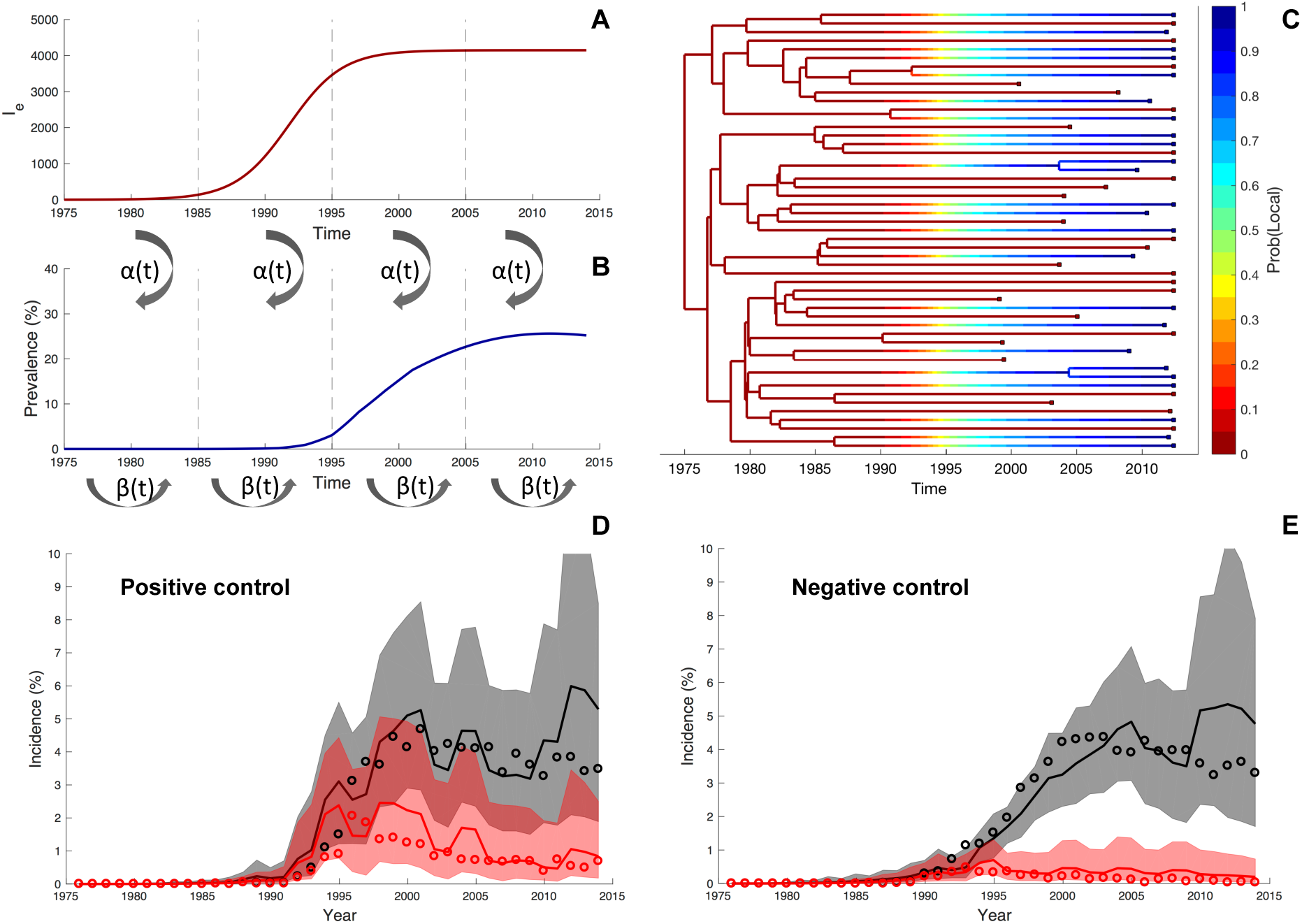
Schematic of the phylodynamic model and its validation on simulated data. Epidemic dynamics simulated under the model showing the number of infected individuals in the external population *I_e_* **(A)** and the local population **(B)**. Transmission events from the external to the local population occurs at rate *α*(*t*) and within the local population at a rate proportional to *β*(*t*). Both of these rates are time-dependent and vary in a piecewise constant manner to accommodate changes in behavior, treatment or other interventions. Although not shown here, viral lineages can also be exported from the local population through transmission to the external population. **(C)** A simulated phylogeny generated under the same phylodynamic model. Here, each lineage is colored according to its probability of being in the local population (blue) based on its sampling location and the estimated transmission rates between populations. **(D-E)** Total incidence (grey) and incidence attributable to external introductions (red) inferred from simulated phylogenies. Solid lines represent the posterior median estimate, shaded regions mark the 95% credible intervals and open circles mark the true yearly incidence known from the simulations. In the positive control **(D)**, we correctly infer that external introductions played a large role in driving and sustaining the local epidemic; whereas in the negative control **(E)** we correctly infer that external introductions only played a minor role in seeding the epidemic.

We therefore developed a new phylodynamic model based on a simple but still realistic epidemiological model using a previously described structured coalescent framework [Volz, 2012] in order to quantify the contribution of external introductions versus local transmission. The model tracks the number of infected individuals in the external population along with local epidemic dynamics (Fig 2A-B). The model also probabilistically tracks how lineages in the tree move between populations based on their sampling location and the estimated transmission rates between populations (Fig 2C). Tracking the movement of lineages in this way allows us to estimate whether new infections were derived from a local or external source. We then validated our model using phylogenies simulated to reflect different epidemic scenarios. In the first scenario, external introductions play a large role in driving and sustaining the local epidemic (positive control). In the second scenario, external introductions only play a minor role in seeding the epidemic (negative control). In both scenarios, we were able to accurately estimate both the overall epidemic dynamics and the incidence attributable to internal and external transmission (Fig 2D-E).

Using the phylodynamic model, we reconstructed epidemic dynamics in the local population from phylogenies reconstructed from the viral samples. As expected, both prevalence and incidence rapidly grew during the 1990s, and then grew more slowly after 2000 (Fig 3A-B). After 2004, independent estimates of prevalence and incidence based on population-based surveillance data are available from the AHRI. Prevalence estimates based on surveillance data were about 10% higher than our phylodynamic estimates, although both methods estimate a similar growth in prevalence (Fig 3A). The estimates of incidence since 2004 are in closer agreement, with both methods returning a median estimate of yearly incidence between 3-4% (Fig 3B).

**Figure 3.**
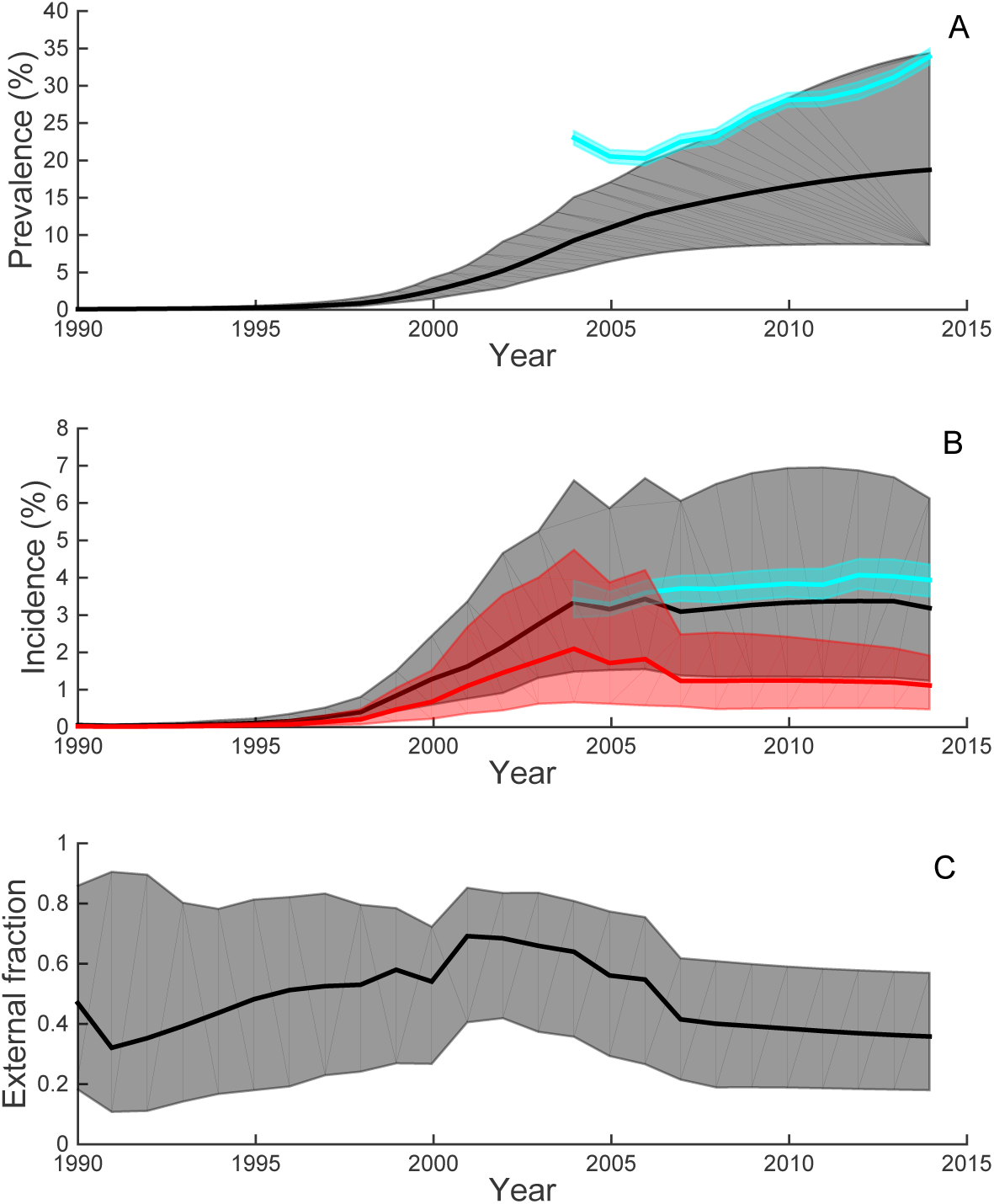
Epidemic dynamics reconstructed from viral phylogenies using the phylodynamic model. **(A)** Prevalence estimates from the phylogeny (grey) and independent surveillance data (blue). **(B)** Total incidence estimated from the phylogeny (grey) and surveillance data (blue). Incidence attributable to external introductions estimated from the phylogeny is shown in red. **(C)** The fraction of incidence attributable to external introductions over time. All solid lines represent the posterior median estimates while shaded regions mark the 95% credible intervals. All estimates represent a posterior average over a set of phylogenies reconstructed from different sub-sampled datasets and thus take into account both phylogenetic uncertainty and sampling variance.

Given that we could reliably reconstruct overall epidemic dynamics from simulated and empirical viral phylogenies, we used the phylodynamic model to quantify the relative contribution of external introductions versus local transmission to overall incidence. During the earliest stages of the local epidemic, the incidence attributable to external introductions was very high (Fig 3B; red). While after 2005 the fraction attributable to external introductions declined (Fig 3B; red), as of 2014 an estimated 35% (95% CI: 20-60%) of all present day infections were due to external introductions (Fig 3C).

## Discussion

Our phylodynamic results on external introductions fit within a growing body of evidence that transmission networks are highly interconnected across communities even at relatively large geographic distances due to human movement patterns. Historians and social scientists have long noted that both mass migration and increased mobility may have played an important role in the rapid spread of HIV through southern Africa [Iliffe, 2006]. In South Africa in particular, historical patterns of circular labor migration among neighboring countries and increased mobility following the end of the Apartheid system may have contributed to the rapid growth of the epidemic [Jochelson et al., 1991, Lurie et al., 1997, Hargrove, 2008]. More recently, phylogenetic studies of HIV in Africa have provided support for frequent viral movement at spatial scales ranging from local communities [Grabowski et al., 2014] to across national borders [Wilkinson et al., 2016]. This frequent movement has been linked to highly mobile individuals such as economic migrants, soldiers, prostitutes and truck drivers [Gray et al., 2009, Wilkinson et al., 2016].

Our phylogenetic and phylodynamic analysis of the HIV epidemic in the Africa Health Research Institute study population revealed that external introductions played and continue to play a vital role in driving the epidemic in rural KZN. A preliminary phylogenetic analysis based on maximum parsimony suggested that a large wave of introductions occurred in the 1990s during the early stages of the South African epidemic. A subsequent analysis using a more realistic phylodynamic model indeed confirmed that the earliest stages of the epidemic were largely driven by external introductions.

Our phylodynamic analysis also suggested that, far from just seeding the local epidemic, external introductions continue to play an important role in sustaining the high incidence of HIV in local KZN populations. This has direct relevance for antiretroviral treatment as prevention (TasP) trials and programs. If most transmission occurs locally, TasP programs should be able to efficiently and cost-effectively reduce incidence [Granich et al., 2009] and could selectively target communities with higher burdens of new infections [Tanser et al., 2009]. However, if most new infections are acquired externally, then increasing local ART coverage may not substantially reduce incidence. Our phylodynamic estimates for one KZN population indicate that the situation likely lies between these two extremes. Our median estimate is that presently 35% (CI: 20-60%) of new infections in the AHRI population are attributable to external introductions, suggesting a substantial number of new infections could be prevented if the source of these infections could also be targeted. Nevertheless, these results imply that the majority of new infection events may be attributable to local transmission, and increasing local ART coverage may still prevent many future infections. However, a recent TasP trial in a population immediately adjacent to the AHRI study area showed no incidence reduction despite increased access to ART [Iwuji et al., 2016]. Identifying the source of new infections will be key to understanding why this trial failed to decrease incidence.

While phylogenies can reveal the movement of viruses between populations, they cannot necessarily reveal where transmission events occurred or how viral lineages first entered a population without additional information about human movement. An external introduction event may result from a visitor transmitting to an individual residing in the local population or while an individual currently residing in the local population was living away or traveling. In our phylodynamic model, we can not distinguish between these two scenarios because they result in identical phylogenetic patterns. The AHRI does however collect data on the residence and migration status of individuals in the study area. This data has shown that prevalence is higher among residents with histories of recent migration [McGrath et al., 2015]. Individuals who spend more time away and travel longer distances outside their community also have a significantly higher risk of acquiring HIV infection [Dobra et al., 2017]. Both of these observations support the idea that local residents may be infected while living or traveling outside of the study area and then migrate or return to the area.

Although phylodynamics is increasingly used to study epidemic dynamics, the validity of estimates derived entirely from viral phylogenies can of course be questioned and have their own limitations. We however feel confident that our estimates accurately reflect the true situation for several reasons. First, we validated our model on simulated phylogenies and found that we could accurately reconstruct total incidence and the fraction of incidence attributable to external introductions in scenarios where introductions both did and did not play a large role in driving epidemic dynamics. Second, we were able to reconstruct dynamics consistent with prevalence and incidence trends estimated from independent surveillance data. Third, while there is no “gold standard” to which we can compare our phylodynamic estimates of external introductions, our estimated fraction of external introductions is consistent with known mobility patterns in the AHRI population where 38% of males and 32% of females are recent migrants or frequently leave the area [Camlin et al., 2010, Muhwava et al., 2013]. Finally, although in this study we focused on quantifying the relative contribution of external introductions to address a long standing question about the role that human mobility plays in local HIV epidemics, preventing future infections ultimately requires a detailed knowledge of where new infections are coming from. Using only phylogenetics to reveal the geographic source of new infections remains a difficult challenge at present. Most large-scale HIV sequencing studies have focused on extensively sampling local populations, while the epidemic remains sparsely sampled at larger spatial scales. Nevertheless, as HIV sequence datasets expand to encompass larger spatial regions, it should be possible to use phylodynamic methods similar to those used here to pinpoint the geographic source of new infections by directly quantifying transmission across multiple communities.

In conclusion, we believe our results demonstrate the power of using phylodynamics to study HIV transmission dynamics in large, generalized African epidemics. In addition to showing that external introductions play an important role in sustaining high HIV incidence in this hyper-epidemic setting in rural KZN, we were also able to estimate population-level incidence with remarkalbe accuracy. We think this also represents a key result as there is currently great interest in using phylodynamics together with large-scale sequencing data to quantify incidence and the effect of interventions without the need for expensive longitudinal cohorts [Dennis et al., 2014, Cohen, 2015]. To this end, a recent simulation study has demonstrated that it may be possible to estimate incidence from phylogenies in the context of African HIV epidemics [Ratmann et al., 2017]. Here, we confirm for the first time using empirical sequence data that accurate estimates of HIV incidence are possible using a relatively simple model that incorporates recent advances in phylodynamics [Volz, 2012, Mueller et al., 2016]. We note in closing that this model is now freely available as an add-on to the widely used phylogenetic software package BEAST 2 [Bouckaert et al., 2014].

## Methods

### Study population

We focused on the Africa Health Research Institute (AHRI) study population in KwaZulu-Natal, South Africa. The AHRI demographic surveillance area (DSA) is located 200km north of Durban with a mostly rural or peri-urban population. Demographic data was compiled from the Africa Centre Demographic Information System (ACDIS) between 2000 and 2015 [Tanser et al., 2008]. There were approximately 87,000 people under surveillance in this population at any given time, with an adult population of males aged 15-54 and females aged 15-49 of 55,000. Prevalence and incidence data were also made available from ongoing surveillance.

### Population-based HIV surveillance

A population-based HIV survey has been performed within the DSA since 2004 and we compared our phylodynamic estimates of prevalence and incidence against data from this survey. Trained field-workers visit households every 12 months and identify eligible participants aged 15 to 49 years. After obtaining consent, the field workers then extract blood according to the UNAIDS and WHO Guidelines for Using HIV Testing Technologies in Surveillance. Of the eligible participants contacted, approximately 60% agree to be tested at least once during the study period. Prevalence was determined from the number of individuals who tested HIV-positive during these yearly visits.

We calculated incidence rates using the surveillance data from the subset of eligible participants who had two or more HIV tests, of which the first was a valid HIV-negative result [Vandormael et al., 2014]. During the 2004 to 2014 surveillance period, we observed 2,557 seroconversion events for the 17,417 participants in the incidence cohort, irrespective of their residency status or time spent in the DSA. Due to periodic HIV testing, however, we did not know the exact dates of the seroconversion events. We therefore used an Monte Carlo based approach to impute a random seroconversion event between the *i*th participant’s latest-negative (*l_i_*) and earliest-positive (*r_i_*) test dates [Vandormael et al., 2017].

### Sequence data and phylogenetic analysis

Partial HIV-1 polymerase (*pol*) sequences were collected as part of population-based surveillance conducted in 2011 and 2014. HIV-1 viral load tests were done on all dried blood spot samples that tested positive by serology (SD Bioline ELISA). Only samples from ART-naive participants with viral loads greater than 10,000 RNA copies/ml were genotyped. More information about the genotyping process is described in Manasa et al. [2016]. In total, 1,068 sequence samples from the DSA were included in our analysis. In addition to these samples, we used a large dataset containing 11,289 unique subtype C *pol* sequences previously described in Wilkinson et al. [2016] as a regional background dataset to help identify external introductions. This dataset contained other sequences from South Africa (n=7739) as well as sequences sampled between 1989 and 2014 from Angola (n=9), Botswana (n=863), DRC (n=25), Malawi (n=352), Mozambique (n=342), Swaziland (n=47), Tanzania (n=168), Zambia (n=1,476) and Zimbabwe (n=268).

Maximum likelihood (ML) phylogenetic trees were reconstructed using FastTree2 [Price et al., 2010]. The ML trees were then dated using Least Squares Dating [To et al., 2015], so that branch lengths were given in units of real calendar time. For dating, we assumed a molecular clock rate of 2.0 × 10^3^, which falls in the center of previously estimated clock rates for subtype C [Wilkinson et al., 2016].

In a preliminary analysis, ML trees were reconstructed from an alignment containing all sequences in the background dataset together with all samples from the AHRI. To identify potential external introductions into the local population, we reconstructed the ancestral location of each internal node in the ML trees using the Fitch parsimony algorithm [Sankoff, 1975]. External introductions were assumed to occur whenever a child node reconstructed to be in the local population had a parent node reconstructed to be in the external population. The height of the child node (in units of time) was then used as a proxy for the probable time of introduction.

For the phylodynamic analysis, phylogenetic trees were reconstructed from all AHRI sequences and an equal number of sequences randomly sampled from the background dataset. This was done to reduce the computational cost of fitting the phylodynamic model. To take into account phylogenetic uncertainty and variability across sub-sampled datasets, the phylodynamic analysis was replicated on 10 phylogenies each reconstructed from a different set of sequences sub-sampled from the full regional background dataset. Estimates provided in the Results represent an average over these 10 phylogenies and sampling replicates (see MCMC details below).

### Phylodynamic model

Our phylodynamic model divides the host population into a local and an external population. Within the local population, transmission dynamics are modeled using a standard SIR epidemiological model where the number of susceptible (*S*), infected (*I*), or removed (*R*) individuals change over time according to the differential equations:

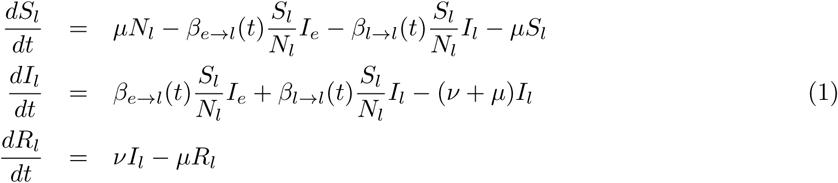

The subscripts denote whether a host resides in the local (*l*) or external (*e*) population. Two sources of transmission contribute to new infections within the local population. The first source, the 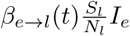 term in (1), represents external transmissions into the local population from the external infected population and corresponds to the *α*(*t*) terms in Fig 2. The second source, the 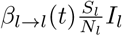 term in (1), represents transmission within the local population and corresponds to the *β*(*t*) terms in Fig 2. All transmission rates *β*(*t*) are allowed to vary over time to accommodate changes in risk behavior, treatment or other non-modeled factors. The birth rate *μ* and removal rate *ν* are assumed to be constant over time.

We simply assume that the external infected population size *I_e_* grows logistically over time according to a SIS model. The number of susceptible (*S_e_*) and infected (*I_e_*) hosts in the external population change over time as follows:

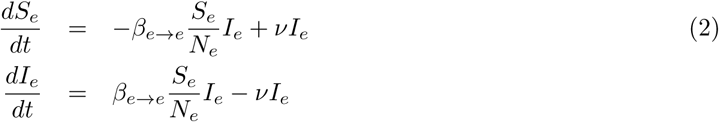

We set the initial infected population size 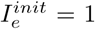 in 1975. While this model ignores the considerable spatiotemporal complexity in HIV dynamics within southern Africa, it captures the main trend in HIV dynamics over time—rapidly increasing growth followed by stabilization in prevalence. We therefore view *I_e_* as the effective number of infected individuals living in the region who could have transmitted to an individual living in the local population.

We assume that the time-dependent transmission rates, *β*_e→l_(*t*) and *β_l_*_→_*_l_*(*t*), are constant within a given time interval, but can change between intervals in a piecewise-constant manner. To obtain a smoothed estimate of how these parameters change over time, we placed a Gaussian Markov Random Field prior on each parameter. Such GMRF models have previously been used in phylodynamics to prevent large fluctuations in parameter values between neighboring time intervals by penalizing against overly large changes [Minin et al., 2008]. The prior probability on a given sequence of time-dependent parameters *γ*_1:_*_T_* is computed as

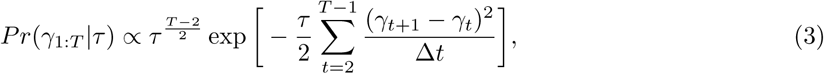
where *τ* is the precision parameter that controls the expected autocorrelation in parameter values between neighboring time intervals. The precision parameter *τ* was inferred separately for each time-dependent parameter. The time interval Δ*t* between change points was fixed at two years between 1990 and 2008. Time-dependent parameters were assumed to be constant prior to 1990 and after 2008 to prevent over-fitting during time periods when the phylogeny was relatively uninformative about changes in epidemic dynamics.

The structured coalescent framework of Volz [2012] was used to compute the likelihood of the reconstructed phylogenies under our phylodynamic model, which has previously been used to analyze HIV transmission dynamics [Volz et al., 2013, Rasmussen et al., 2014]. The likelihood of a given phylogeny under a general structured coalescent model is:

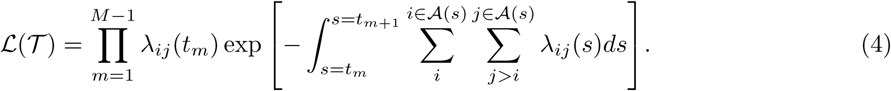

For a tree containing *M* samples, the total likelihood is the product of the likelihood of each of the *M −* 1 coalescent events and the waiting times between events. The likelihood depends on the pairwise coalescent rate λ*_ij_* (*t*) at which two lineages *i* and *j* coalesce at time *t*. The total coalescent rate is then computed by summing over all pairs of lineages in the set of lineages ***A***(*s*) present in the phylogeny at time s, which is allowed to change within coalescent intervals due to sampling.

As shown in Volz [2012], under a structured epidemiological model with more than one type of infected individual, the pairwise coalescent rate is:

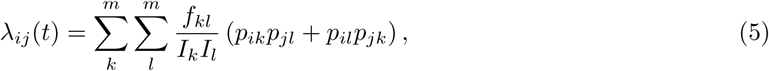
where *p_ik_* is the probability that lineage *i* is in state *k* and *p_jl_* is the probability that lineage *j* is in state *l*. *I_k_* and *I_l_* are the number of infected individuals in populations *k* and *l*, respectively.

The lineage state probabilities can be computed using a system of differential equations that describe how the probability of lineage *i* being in state *k* evolves backwards in time based on its sampling location:

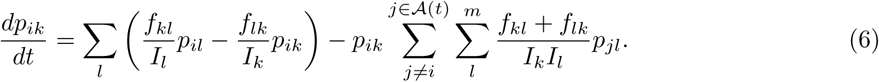

Here we have modified the equations originally introduced in Volz [2012] by adding the final term in (6), which takes into account how the relative probability of lineage *i* being in state *k* changes conditional on the observation that the lineage has not coalesced with any other lineage *j* in the phylogeny. This was shown by Mueller et al. [2016] to improve parameter estimates under asymmetric population dynamics and sampling.

Under our two population HIV model, *f_kl_* represents the transmission rate between populations *k* and *l*. In matrix notation, we have the transmission rate matrix:

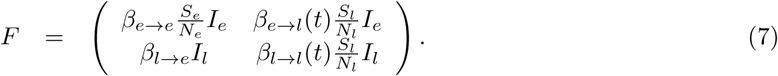

Transmission from the local to the external population allows lineages in the local population to move back into the external population, but the rate *β_l_*_→_*_e_* is assumed to be constant over time. Transmission in this direction does not affect the number of infected individuals in the external population, as this is assumed to be much larger than the number of cases exported from the local populations.

We implemented this phylodynamic model in BEAST 2 [Bouckaert et al., 2014] as an add-on package named Marula. Source code and input XML files that can be used to replicate our analysis are freely available at https://github.com/davidrasm/Marula. The posterior distribution of all model parameters and epidemic dynamics were inferred using BEAST’s built-in MCMC sampler. Due to the large size of the phylogenies, replicate MCMC runs were performed on 10 different fixed ML phylogenies reconstructed from different sub-sampled sequence datasets (see above) rather than jointly estimating the full phylogeny while simultaneously fitting the phylodynamic model. Samples from each MCMC replicate were then pooled to obtain posterior estimates averaged over the different ML phylogenies. Each MCMC replicate was run for at least one million iterations.

For inference, weakly informative priors were placed on all estimated parameters (Table 1). In addition, a few demographic parameters were assumed to be known from the AHRI DIS (Table 2). The local population size *N_l_* reflects the adult population size of males 15-54 and females 15-49 years old. The population was assumed to be at demographic equilibrium with a constant birth/death rate of 2.8% per capita each year.

**Table 1.** Prior distributions on all estimated parameters.

| Parameter | Name | Prior dist. | Prior value |
| --- | --- | --- | --- |
| $\beta_{l \rightarrow l}$ | Local transmission rate | LogNorm | $\mu = -2.3026$; $\sigma = 1.0$ |
| $\beta_{e \rightarrow e}$ | External transmission rate | LogNorm | $\mu = -0.5065$; $\sigma = 1.0$ |
| $\beta_{e \rightarrow l}$ | External introduction rate | LogNorm | $\mu = -2.3026$; $\sigma = 1.0$ |
| $\beta_{l \rightarrow e}$ | Local export rate | LogNorm | $\mu = -2.3026$; $\sigma = 1.0$ |
| $N_e$ | External pop size | Uniform | min= 0.0; max= 250,000 |
| $\nu$ | Removal rate | LogNorm | $\mu = -2.2769$; $\sigma = 0.1$ |
| $\tau$ | GMRF precision | Gamma | $\alpha = 0.01$; $\beta = 0.01$ |

**Table 2.** Demographic parameters fixed at constant values.

| Parameter | Name | Value |
| --- | --- | --- |
| $N_l$ | Local pop size | 55,000 |
| $\mu$ | Birth/death rate | 2.8% yr<sup>-1</sup> |

## Acknowledgments

We would like to thank Andrew Tomita for providing help and data from the Africa Centre Demographic Information Service. D.A.R. was funded by the ETH Zürich Postdoctoral Fellowship Program, the Marie Curie Actions for People COFUND, and a travel fellowship from the Wellcome Trust to visit the AHRI. TdO, EW, AV are supported by a research Flagship grant from the South African Medical Research Council (MRC-RFA-UFSP-01-2013/UKZN HIVEPI), a Royal Society Newton Advanced Fellowship (TdO), and the European Union?s Horizon 2020 Research and Innovation Programme under grant agreement no 634650. DP and AHRI are funded by the Wellcome Trust.

